# Keratinocyte-Derived Exosomes in Painful Diabetic Neuropathy

**DOI:** 10.1101/2024.08.21.608803

**Authors:** James Coy-Dibley, Nirupa D. Jayaraj, Dongjun Ren, Paola Pacifico, Abdelhak Belmadani, Yi-Zhi Wang, Kamil K. Gebis, Jeffrey N. Savas, Amy S. Paller, Richard J. Miller, Daniela M. Menichella

## Abstract

Painful diabetic neuropathy (PDN) is a challenging complication of diabetes with patients experiencing a painful and burning sensation in their extremities. Existing treatments provide limited relief without addressing the underlying mechanisms of the disease. PDN involves the gradual degeneration of nerve fibers in the skin. Keratinocytes, the most abundant epidermal cell type, are closely positioned to cutaneous nerve terminals, suggesting the possibility of bi-directional communication. Exosomes are small extracellular vesicles released from many cell types that mediate cell to cell communication. The role of keratinocyte-derived exosomes (KDEs) in influencing signaling between the skin and cutaneous nerve terminals and their contribution to the genesis of PDN has not been explored. In this study, we characterized KDEs in a well-established high-fat diet (HFD) mouse model of PDN using primary adult mouse keratinocyte cultures. We obtained highly enriched KDEs through size exclusion chromatography and then analyzed their molecular cargo using proteomic analysis and small RNA sequencing. We found significant differences in the protein and microRNA content of HFD KDEs compared to KDEs obtained from control mice on a regular diet (RD), including pathways involved in axon guidance and synaptic transmission. Additionally, using an *in vivo* conditional extracellular vesicle (EV) reporter mouse model, we demonstrated that epidermal-originating GFP-tagged KDEs are retrogradely trafficked into the DRG neuron cell body. Overall, our study presents a potential novel mode of communication between keratinocytes and DRG neurons in the skin, revealing a possible role for KDEs in contributing to the axonal degeneration that underlies neuropathic pain in PDN. Moreover, this study presents potential therapeutic targets in the skin for developing more effective, disease-modifying, and better-tolerated topical interventions for patients suffering from PDN, one of the most common and untreatable peripheral neuropathies.

## INTRODUCTION

Diabetes mellitus affects 29.3 million adults with pre-diabetes in an additional 115.9 million^1^. Painful diabetic neuropathy (PDN) is a disabling, intractable, and common syndrome occurring in approximately 25% of diabetics^2–5^. The associated neuropathic pain significantly impacts the quality of life for patients^6^. Despite the high prevalence and impact, current therapies for PDN have limited effects in treating pain^7–10^, fail to remediate the damage to nerves, and have side effects associated with their systemic administration^10–12^. Therefore, there is an urgent need for better tolerated and more effective therapies for PDN.

PDN is characterized by neuropathic pain associated with dorsal root ganglion (DRG) nociceptor hyperexcitability and the degeneration with loss or retraction of the cutaneous DRG neuron axons that innervate the skin (small fiber neuropathy)^13,14^. A critical barrier to developing effective treatments for PDN is the lack of understanding of the molecular mechanisms leading to neuropathic pain and small fiber neuropathy.

Keratinocytes are the most abundant epidermal cell type. Recent studies have discovered a new role for keratinocytes in mediating innocuous and noxious touch and thermal sensation in healthy skin^15^. Keratinocytes detect touch stimuli in the skin and transmit mechanical information related to pressure and brushing^16,17^. Optogenetic inhibition of keratinocytes *in vivo* inhibits the responses to noxious mechanical and thermal stimuli^17,18^. There is also evidence that keratinocytes may contribute to persistent neuropathic pain. A study involving the transplantation of human keratinocytes into rodents with transected nerves showed increased excitability of DRG neurons and chronic pain *in vivo*^19^. However, the specific role of keratinocytes in PDN has not been widely investigated.

The skin is a highly complex biological system. Along with nociceptive DRG neurons, various other neuronal subpopulations terminate in the skin, both in the dermis and the epidermis^5^. Keratinocytes are closely juxtaposed to cutaneous nerve terminals, suggesting that there may be bidirectional communication. Interestingly, in rodents and human skin, cutaneous nerve terminals in the epidermis form synapse-like contacts and also tunnel through keratinocytes, where they form connexin 43-positive gap junctions, enabling direct cellular communication^20,21^. However, the functional implications of such observations remain unclear. One such ubiquitous mode of intercellular communication recently garnering more appreciation in the skin is mediated by extracellular vesicles (EVs).

Exosomes, which are small EVs composed of lipids, proteins and nucleic acids^22–25^, were initially posited to be involved in removing cellular waste^26^. However, a substantial body of research now suggests a wider role in intercellular communication. Exosomes are released from most cell types and have been linked to several neurogenerative diseases^27,28^ and the progression of different cancers^29^. Keratinocyte-derived exosomes (KDEs) have demonstrated their ability to modulate melanocyte pigmentation^30^, regulate dermal fibroblast gene expression^31^, mediate crosstalk with macrophages in cutaneous wound healing^32^, and play a crucial role in dermal immune responses in psoriasis ^33,34^. Additionally, EVs derived from mesenchymal stem cells can directly alter the excitability of DRG nociceptors in mice^35^. Exploration of the role of exosomes in diabetes, however, has primarily focused on adipose tissue and inter-organ communication^36–38^. Recent studies have unveiled a potential role for exosomes in impaired wound healing associated with diabetes^39^. Yet, research on the effects of exosomes on diabetic neuropathy is limited and has been conducted using exosomes isolated from mesenchymal^40,41^ or Schwann cells^42^. Notably, exosomes isolated from mesenchymal stromal cells have shown promise in ameliorating peripheral neuropathy in a mouse model of diabetes^40^. Conversely, exosomes derived from high glucose-stimulated Schwann cells have been found to promote the development of diabetic neuropathy in mice^42^. A rigorous and comprehensive investigation of keratinocyte-derived exosomes and their role in PDN represents an important, yet understudied frontier in pain and peripheral neuropathy research.

Using size exclusion chromatography, we obtained enriched keratinocyte-derived exosomes from mice and performed an unbiased molecular cargo characterization with proteomics and small RNA sequencing. We found significantly altered protein and microRNA content of keratinocyte-derived exosomes from high-fat diet (HFD)-induced PDN compared to regular diet (RD) control mice. Altered pathways were involved in axon guidance and synaptic transmission. Additionally, using an *in vivo* conditional EV reporter mouse line, we demonstrated that epidermal-originating GFP-tagged keratinocyte-derived exosomes are retrogradely trafficked into the DRG neuron cell body. Overall, we present evidence that supports keratinocyte-derived exosomes as a novel interaction pathway between epidermal keratinocytes and DRG neurons and that altered cutaneous EV-trafficking may play a functional role in the development of the small fiber neuropathy observed in PDN.

## RESULTS

### Keratinocytes release a diverse population of exosomes

To study keratinocyte-derived exosomes (KDEs), we fractionated cell-conditioned medium (CCM) from cultured adult mouse primary keratinocytes^17,18^ using a qEV 35-nm size-exclusion chromatography column^43^ (Figure 1A). We observed an increasing total protein concentration with each successive fraction (Figure 1B; silver stain) and immunoblotted for known exosome markers. Fractions 2 and 3 were enriched with the exosome-associated cargo markers Alix, Tsg101, and Syntenin-1 as well as the transmembrane tetraspanins CD63 and CD81^23–25,44^. These fractions were devoid of GM130 and Calnexin, which are Golgi-associated proteins^45^ (Figure 1B). Thus, pooled fractions 2 and 3, representing our highly enriched KDE fractions with minimal free-floating protein contamination, were used for all subsequent analyses^46^.

**Figure 1:**
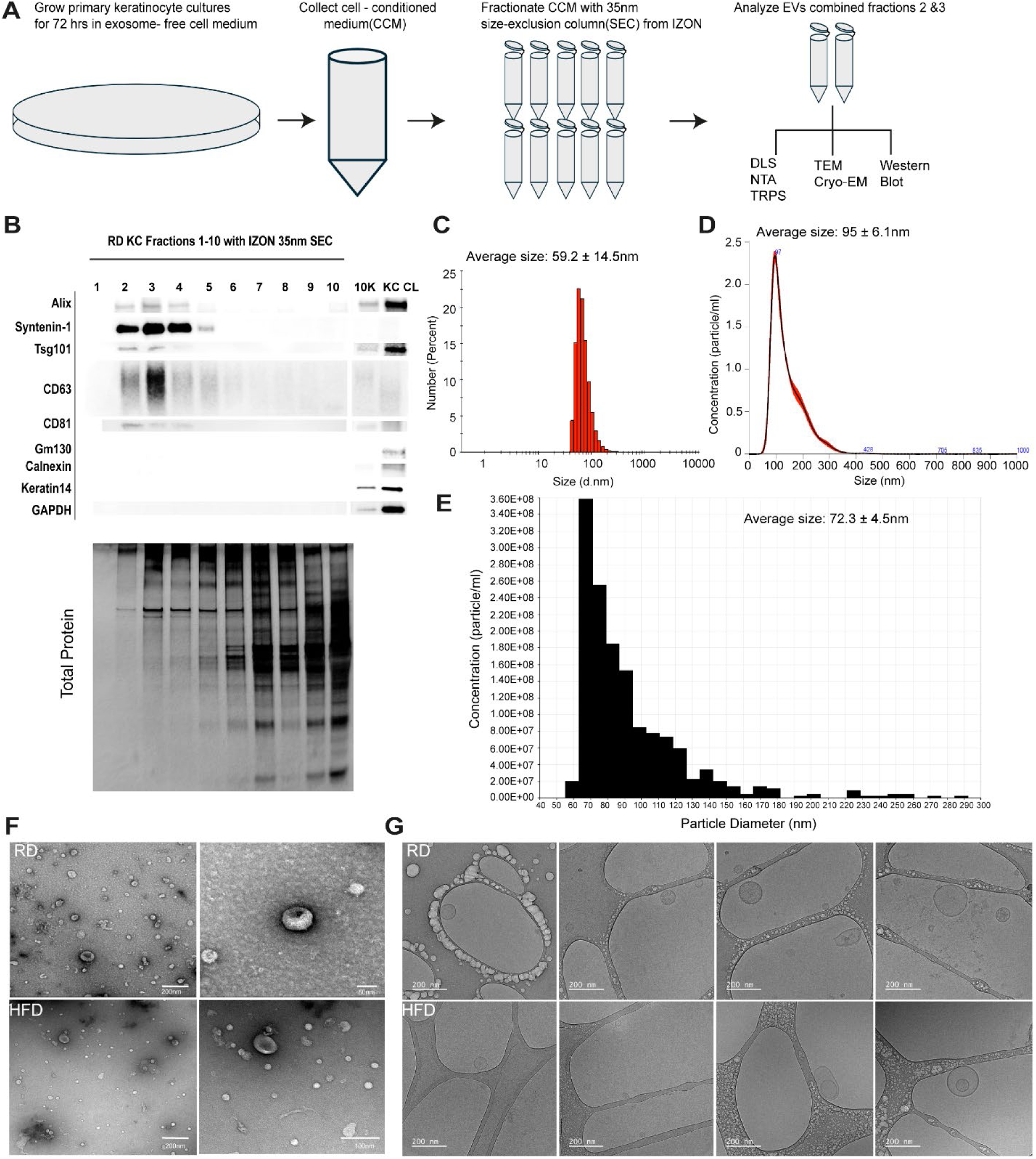
Keratinocytes release a diverse population of exosomes. **A)** Workflow schematic for keratinocyte-derived exosome (KDE) isolation using size exclusion chromatography (SEC). **B)** Immunoblotting the ten SEC fractions revealed that exosome-associated proteins ALIX, Syntenin, Tsg101, CD63, and CD81 were enriched in fractions 2 and 3 relative to the total protein concentration (silver stain). **C)** KDEs analyzed with dynamic light scattering had a size range of 59.2 ± 14.5nm (mean ± StDev). N=7 across biological replicates from both male and female mice. **D)** KDEs analyzed using nanoparticle tracking analysis produced a concentration peak particle size of 95 ± 6.1 nm (mean ± StDev). N=2 male biological replicates. **E)** KDEs analyzed using tunable resistive pulse sensing (TRPS) determined that the mean size for RD KDEs was 97.7 ± 4.5nm, with a concentration peak of 72.3 ± 4.5nm (mean ± StDev). N=2 biological replicates. **F)** Nanovesicles from combined fraction 2/3 were visualized with negative stain EM. N=11 across 5 biological replicates from male and female mice. **G)** Combined SEC fractions 2/3 were visualized with cryo-EM, demonstrating several diverse populations. N=2 biological replicates, one male and one female.

To investigate the role of KDEs in painful diabetic neuropathy, we employed the clinically relevant and well-established HFD model of PDN^47–50^, developed by feeding mice a high fat diet for ten weeks, leading to obesity, glucose intolerance and mechanical allodynia with small-fiber degeneration^48–50^. Cultured keratinocytes from the RD and HFD mice were grown to 90% confluency in low-calcium, EV-depleted medium for scratch assays. The HFD keratinocytes had impaired wound healing during the 3 days of observation (Supplemental Figure 1A-B), consistent with what has been shown in scratch assays with high glucose exposure^51^ and what is observed in diabetic patients^52^. Fraction 2/3 KDEs had sizes within the range of exosomes of 59.2 ± 14.5nm (RD; Figure 1C) and 70 ± 26.3nm (HFD; Supplemental Figure 2B) by dynamic light scattering (DLS) and concentration peaks at 95 ± 6.1nm (RD; Figure 1D) and 86 ± 8.7nm (HFD; Supplemental Figure 2C) by nanoparticle tracking analysis (NTA). Tunable resistive pulse sensing (TRPS) determined that the mean size average for RD KDEs were 97.7 ± 4.5nm with a concentration peak of 72.3 ± 4.5nm (Figure 1E). KDEs from both RD and HFD were visualized via negative staining (Figure 1F) and cryo-electron microscopy (Figure 1G), which revealed the expected crescent morphology and intact vesicular structure.

### Keratinocytes release soluble factors that encourage DRG neuron axonal growth

Given that loss of cutaneous innervation is reported in PDN^13,14,49^, we next sought to investigate the role of KDEs on DRG axonal growth. We employed a microfluidics co-culture system with primary adult mouse DRG neurons in one chamber and adult mouse keratinocytes in the other (Figure 2A). This setup allowed for cell medium exchange only through the microchannels between the chambers. Interestingly, we found that DRG neuronal axons crossed the microchannels separating the two chambers at a higher rate when grown in a co-culture with keratinocytes compared to when grown alone, indicating that keratinocytes release soluble growth factors that promote axonal growth (Figure 2B-C).

**Figure 2:**
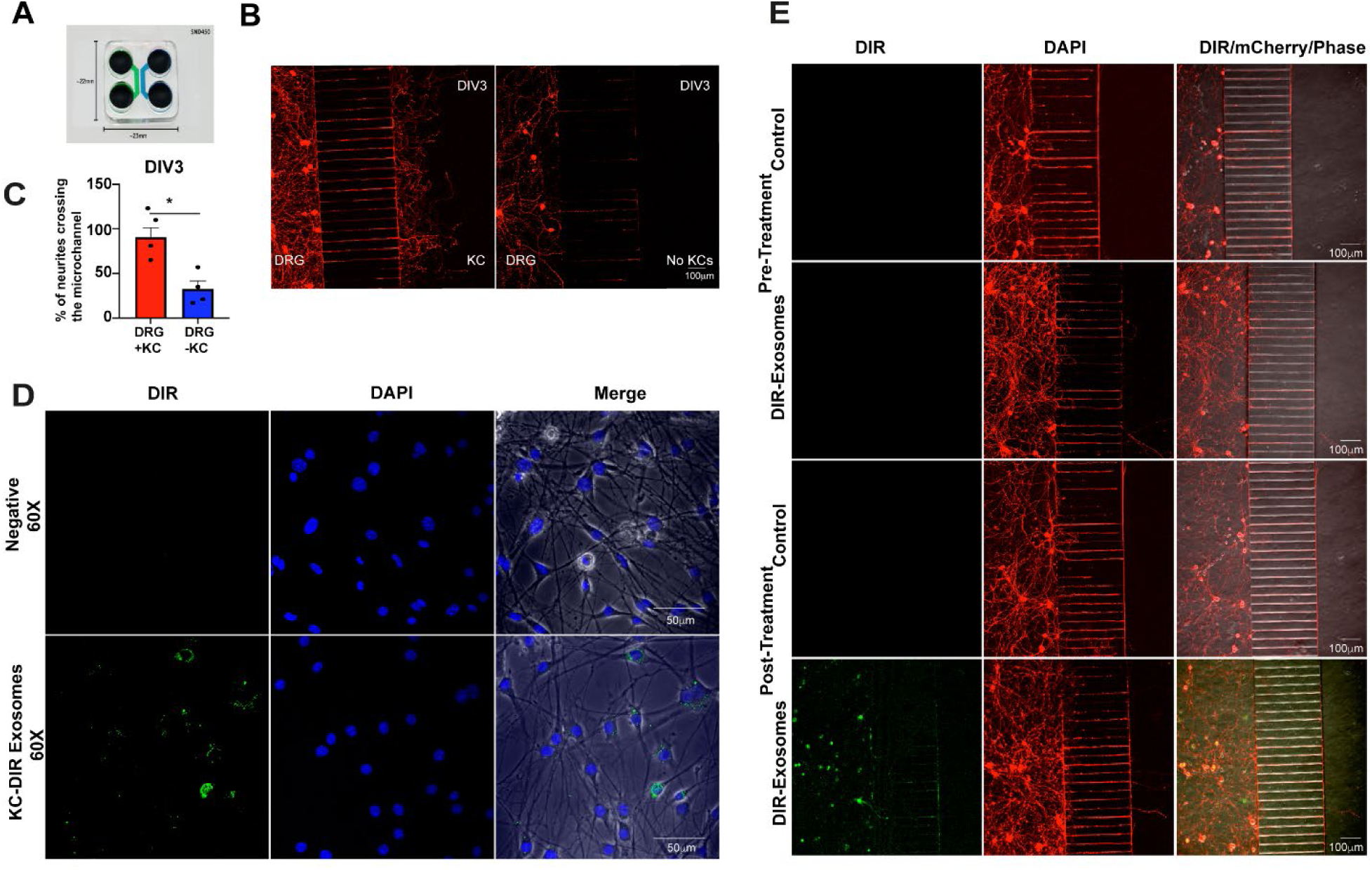
Isolated exosomes from SEC fraction 2/3 are functionally retrogradely trafficked by DRG neurons *in vitro*. **A)** Representative picture of microfluidics device. **B)** The presence of keratinocytes co-cultured with DRG neurons in a microfluidic paradigm encouraged neurite outgrowth crossing through the microchannels connecting both chambers. **C)** We observed a significant increase in the neurite microchannel crossing when co-cultured in the presence of keratinocytes (p ≤ 0.05, one-tailed, paired t-test; n=5 with KCs and n=4 without KCs paired with DRG primary cultures). **D)** KDEs from SEC Fr2/3 are functionally internalized by primary DRG neurons. N=4 across 2 biological replicates. **E)** KDEs are retrogradely trafficked into the neuron cell body through the neurites in a microfluidic paradigm. N=4 biological replicates.

To test whether mouse KDEs are functional, we labeled them with DIR, which is a carbocyanine DiOC18(7) lipophilic fluorescent dye, and added them to primary DRG neuron cultures. We observed a robust uptake of these KDEs by both the cell bodies and neurites of the neurons (Figure 2D). To better model physiological relevance, we then cultured DRG neurons in one chamber of our microfluidic system and allowed their neurites to occupy all the microchannels, thus preventing the free flow of medium between chambers. We then added DIR-labeled KDEs to the empty chamber to test whether they could be transported through the neurites to the DRG neuron cell body. Indeed, DIR-labeled KDEs were readily detected in the DRG neuron cell bodies 16 hours post application, suggesting the retrograde transport of DIR-labeled KDEs through the neurites^53^ (Figure 2E).

### Keratinocyte-derived exosomes alter their protein cargo in painful diabetic neuropathy

We characterized the KDE protein content in the context of PDN using a proteomic approach^54^. The analysis was performed on the pooled fractions 2/3 using liquid chromatography-tandem mass spectrometry (LC-MS/MS) for both RD and HFD KDEs (Figure 3A). A gene ontology enrichment analysis on proteins detected in pooled fractions 2/3 for both groups clustered in EV categories, suggesting a robust small EV enrichment for both RD and HFD KDEs (Figure 3B). Importantly, we found similar quantities based on the number of spectral counts of the canonical exosome-associated proteins Alix, Tsg101, and Syntenin-1 in both samples (Figure 3C). Fractions 2/3-enriched proteins also significantly clustered in the GO enrichment term ‘Axon Development,’ (Figure 3C; Insert), suggesting neuron-keratinocyte communication via exosomes. Moreover, we identified 90 differentially expressed exosome-associated proteins (EAPs; FC ≥ 1.5, p < 0.05) between RD and HFD KDEs, with biological replicates clustering by group (Figure 3D-E). The differentially expressed REACTOME pathway with ‘MAPK Family Signaling Cascades’ included both Mapk1 and Mapk3 (Figure 3G left; Adjusted p-value ≤ 0.05), with Mapk1 found in fraction 2/3 (Figure 3F). Furthermore, a gene ontology enrichment analysis on the EAPs revealed ‘Wound Healing’ as differentially expressed in HFD KDEs (3G middle; Adjusted p-value ≤ 0.05), consistent with the known impairment in diabetic patients^55^ and as shown in our 2D primary HFD keratinocyte scratch assays (Supplemental Figure 1A-B). Notably, several annexins were differentially expressed, with annexin VII in fraction 2/3 of our immunoblots (Figure 3F); annexins play a role in ESCRT-III mediated plasma membrane repair^56–59^ and wound healing^56^. Another differentially expressed pathway was ‘Neurotransmitter Secretion’ (Figure 3G right; adjusted p-value ≤ 0.05), consistent with the physical localization of DRG nerve afferents, which tunnel through the keratinocyte cytoplasm to form synapse-like contacts^20,21^. Taken together, these differentially expressed GO pathways support an alteration in the neuron-keratinocyte communication pathways in diabetes-related PDN.

**Figure 3:**
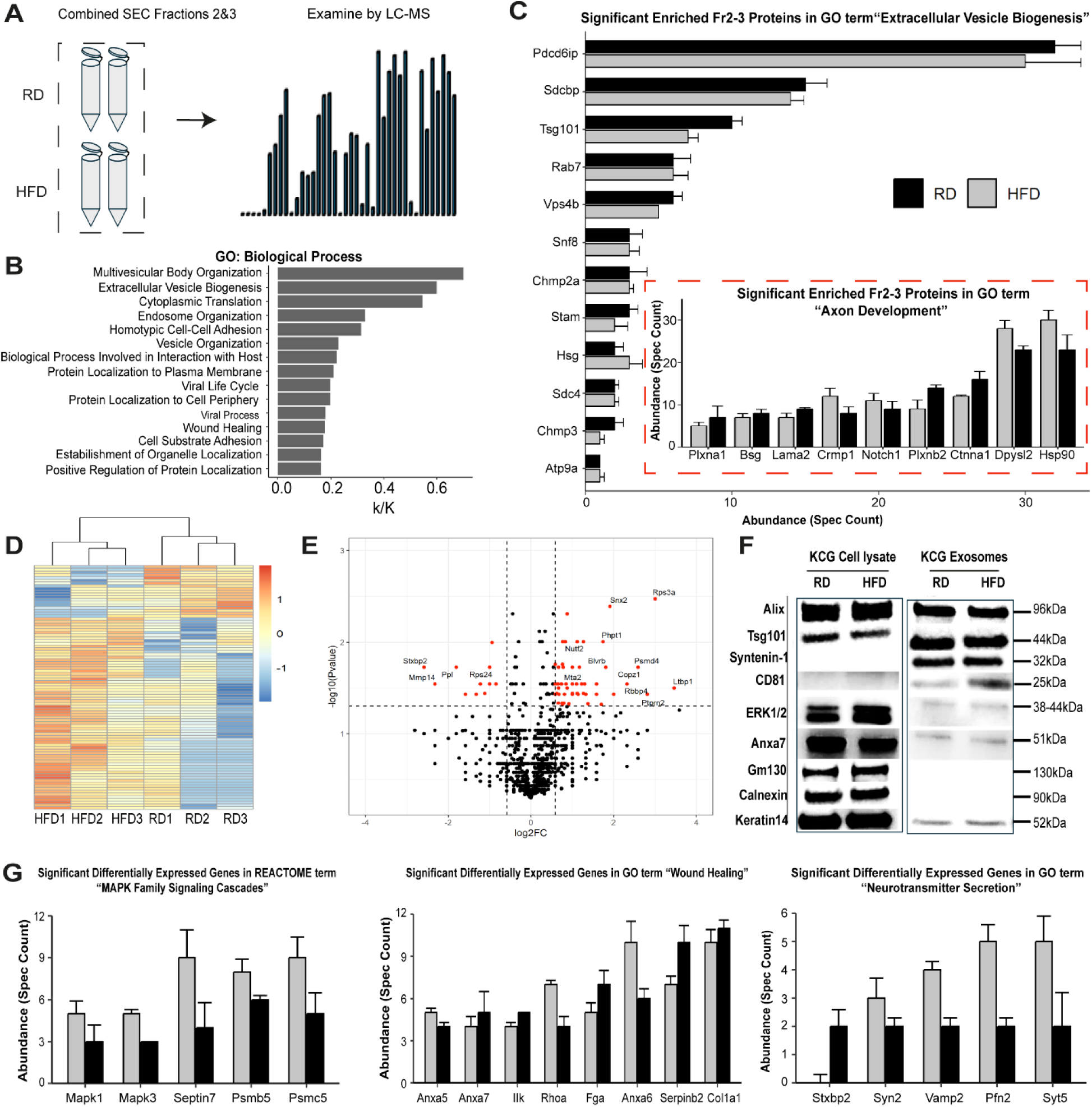
Keratinocyte-derived exosomes (KDEs) significantly alter their protein cargo in painful diabetic neuropathy. **A)** Workflow schematic depicting exosome proteomic analysis. N=3 biological replicates for each group from male mice. **B)** The top pathways from a gene ontology enrichment analysis of proteins present in each of the three biological replicates in both experimental groups suggested EV enrichment. Bonferroni adjusted p-value ≤ 0.01. **C)** Select panel of proteins associated with GO Term ‘EV Biogenesis’ revealed similar spectral counts for exosome markers Alix, Syntenin-1, and Tsg101 between both groups. Insert: GO Term ‘Axon Development’ revealed several unverified notable proteins as exosome cargo. Mean ± SEM. GO Terms Bonferroni adjusted p-value ≤ 0.01. **D)** There were 90 significant differentially expressed exosome-associated proteins (EAPs; FC ≥ 1.5 and paired t-test, one-way, p ≤ 0.05) between RD and HFD KDEs that clustered by group. FC calculated as average HFD/RD spectral count for each protein. **E)** Representative volcano plot with 90 EAPs with FC ≥ 1.5 and p ≤ 0.05 (labelled red). There were 11 downregulated and 79 upregulated EAPs. **F)** Western blot confirms two EAPs, Mapk1 (via ERK1/2 expression) and annexin-7 in SEC Fr2/3. n=3 biological replicates for each group. **G)** Abundances for EAPS in the differentially expressed GO term ‘Wound Healing,’ ‘Neurotransmitter Secretion,’ and the REACTOME term ‘MAPK Family Signaling Cascades.’ Mean ± SEM. Bonferroni adjusted p-value ≤ 0.01.

### The microRNA cargo of keratinocyte-derived exosomes is altered in painful diabetic neuropathy

Exosomes contain gene-modifying RNAs, with microRNAs being the most abundant RNA species^43^. Several microRNAs have been associated with pain in diabetic neuropathy^60^, but the KDE microRNA content has not yet been identified in the context of PDN. Hence, we next investigated the microRNA cargo of RD and HFD KDEs using an unbiased small-library RNA sequencing approach.

The top ten most abundant microRNAs identified, including their variants, accounted for 74.8% of the total small RNA sequenced for both groups (Figure 4A). Additionally, the top two hits, the let-7 and miR-23 families, accounted for almost 40% of the total RNA sequenced for both groups. By cross-referencing 3 separate databases (Figure 2B), the predicted protein targets of these small RNAs were used to run a KEGG enrichment analysis for let-7 and miR-23, with both revealing ‘Axon Guidance’ as a predicted target pathway (Figure 4C). Between the RD and HFD KDEs, there were 35 significant differentially expressed microRNAs that clustered by group (Figure 4D) and 33 with a fold change (FC) ≥ 1.5 (Figure 4E). By cross-referencing the same three databases, the predicted target proteins from each microRNA were run through a gene ontology enrichment analysis with three producing significant target pathways (Figure 4F; Supplemental Figure 3F-G). Both miR-684, which has been implicated in multiple sclerosis^61^, and miR24-3p, which has been studied in the context of cancers^62,63^ along with its regulation of proliferation related to annexin-6 activity^64^, revealed predicted target axon-guidance related pathways (Figure 4G), further providing compelling evidence for a neuron-keratinocyte communication pathway via KDEs under both normal physiology and PDN. Given that both the proteomic and small RNA sequencing datasets revealed predicted target pathways involved in axon guidance and that painful diabetic neuropathy is accompanied by peripheral nerve degeneration^49^, we next designed an *in vivo* animal model to investigate the interaction between KDEs and DRG neuron afferent fibers.

**Figure 4:**
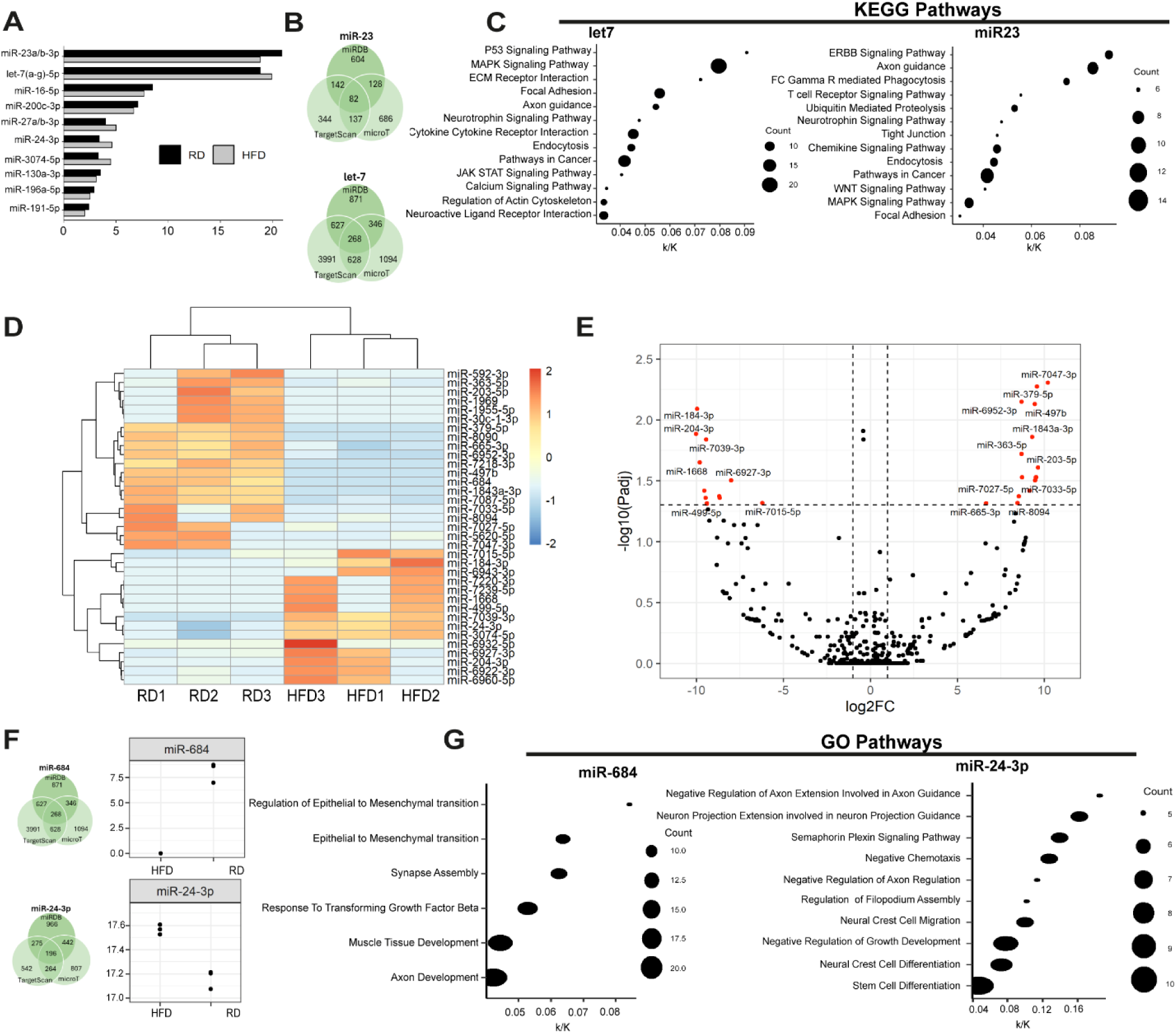
Keratinocyte-derived exosomes (KDEs) alter their small RNA cargo in painful diabetic neuropathy. N=3 biological replicates for male mice in each diet group. **A)** The top ten small RNAs identified in both RD and HFD KDEs represent 74.8% of the total small RNA sequenced. The top two small RNAs, the let-7 and microRNA-23 families, accounted for almost 40% of the total small RNA sequenced. **B)** The predicted protein targets for let-7 and miR-23 were obtained by cross-referencing 3 separate databases: MirDB, TargetScan, and DIANA-microT. The protein targets predicted by all three programs were used for downstream analysis. **C)** The enrichment analysis of the predicted protein targets of let-7 and miR-23 both revealed predicted KEGG pathways related to ‘Axon Guidance,’ suggesting a possible role of KDE small RNAs on keratinocyte-to-neuron communication. KEGG pathways Bonferroni adjusted p-value ≤ 0.01. **D)** There were 35 differentially expressed microRNAs, which clustered by group. Bonferroni adjusted p-value ≤ 0.05. **E)** Of the 33 differentially expressed microRNAs (FC ≥ 1.5, adjusted p-value ≤ 0.05) between RD and HFD, 22 were overexpressed in HFD. FC is defined as HFDavg/RDavg microRNA counts for each small RNA. Bonferroni adjusted p-value. **F)** miR-684 was downregulated in HFD while miR-24-3p was upregulated. Using the same three databases as Figure 4B, predicted target proteins lists were obtained for each. **G)** The gene ontology enrichment analysis for these predicted proteins presented several interesting GO Terms. Both differentially expressed microRNAs predicted axon-related pathways, further providing evidence for an altered keratinocyte-to-neuron communication via exosome cargo. GO Terms Bonferroni adjusted p-value ≤ 0.01.

### Epidermal keratinocyte-derived exosomes are fluorescently labeled with CD63-emGFP

Fraction 2/3 from our keratinocyte CCM consistently immunoblotted for the transmembrane tetraspanins CD63 and CD81^24^ (Figure 1B), in addition to the luminal exosome markers Alix, Syntenin-1, and Tsg101. We crossed the commercially available CD63-emGFP fl/fl mouse line (JacksonLaboratory Strain#:036865) and crossed it with K14-Cre mice to generate a K14-CD63-emGFP EV reporter mouse model (Figure 5A) to label exosomes originating from basal layer keratinocytes and their progeny. We observed robust GFP expression throughout the keratinocyte progeny, with K14 staining co-localizing in basal layer keratinocytes, confirming the limitation of expression to the epidermis (Figure 5B). Furthermore, we immunoblotted whole epidermis cell lysate and detected both the membrane-bound CD63-GFP fusion protein and soluble GFP (Figure 5C), presumably due to endogenous protein recycling. We next cultured keratinocytes from this EV-reporter mouse. As expected, we observed a strong GFP signal along the outer membrane of these keratinocytes (Figure 5D). We next immunoblotted all ten fractions obtained from CCM of primary keratinocyte cultures for GFP and detected a strong GFP signal, both the fusion and soluble forms, in fractions 2/3, which corresponded to our KDE enriched fractions (Figure 5E). Furthermore, isolated GFP-labeled KDEs were functionally internalized by DRG neurons with GFP detected in both the neuron cell bodies and neurites (Supplemental Figure 4A). Given the gene ontology enrichment pathways highlighted from our proteomic and small RNA sequencing experiments, we next sought to better understand the neuron-keratinocyte communication pathway using our EV reporter mouse line.

**Figure 5:**
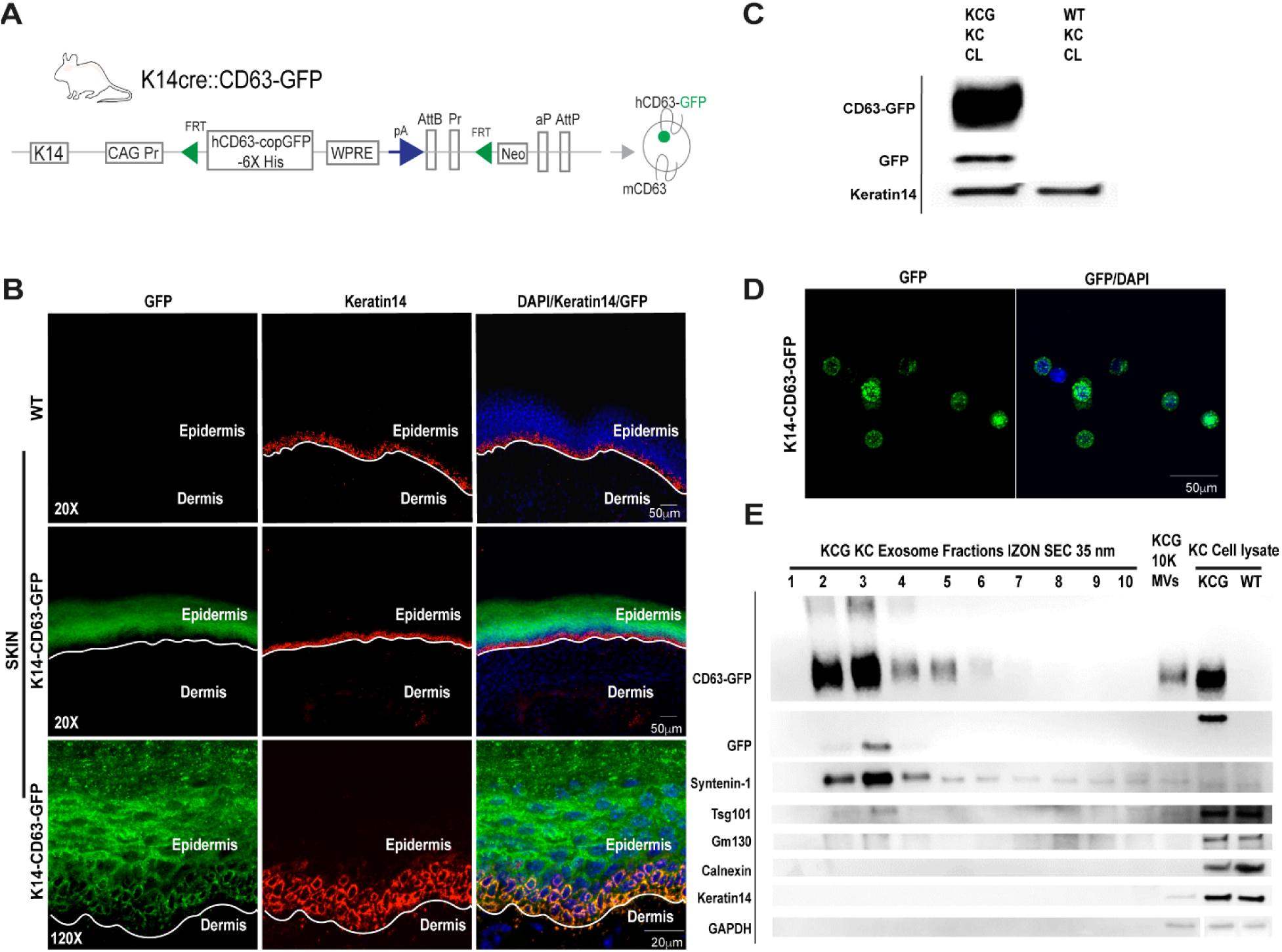
Epidermal keratinocyte-derived exosomes (KDEs) are enriched with GFP in an EV-reporter mouse model. **A)** We created an EV-reporter mouse line by crossing the commercially available CD63-GFP fl/fl mouse line with K14-Cre to create K14-CD63-GFP (KCG) mice. **B)** IHC on cryo-sections of glabrous mouse skin demonstrated GFP expression in all layers of the epidermis with no detectable GFP in the dermis. The GFP (green) signal also co-localized with K14 staining (red), representing the basal layer of the epidermis. **C)** Immunoblotting revealed robust GFP signal in epidermal cell lysate from our EV-reporter mouse line. **D)** GFP was detected in the primary keratinocyte cultures from our EV-reporter mouse line. **E)** Cell-conditioned medium from KCG keratinocyte cultures was run through IZON 35nm SEC columns. Fraction 2/3 was enriched with CD63-GFP and soluble GFP, supporting that KDEs from primary cultured keratinocytes from our EV-reporter mouse line are tagged with the CD63-GFP fusion protein.

### Epidermal keratinocyte-derived exosomes are retrogradely transported to the DRG neuron cell bodies in male and female mice

We further investigated the *in vivo* conditional EV-reporter mouse, shifting our focus towards the DRGs. Notably, we detected a GFP signal in DRG cross-sections from both RD and HFD EV-reporter mice (Figure 6A; Supplemental Figure 4B) and not in WT DRG controls (Figure 6B). Additionally, the cell lysate of K14-CD63-GFP (KCG) DRG neurons immunoblotted for GFP (Figure 6C), further supporting the evidence for the retrograde transport of GFP-positive epidermal exosomes to the DRG neuron cell bodies. We next cultured primary DRG neurons from the EV-reporter mouse model for both RD and HFD. We observed robust GFP expression in both groups (Figure 6D) with no GFP signal in WT controls (Figure 6E). The GFP observed in these primary DRG neuron cultures presented in the same punctate pattern observed in the DIR-labeled exosomes (Figure 2D) and GFP-labeled exosomes applied to primary DRG cultures^46^ (Supplemental Figure 4A). Indeed, we detected these puncta in both the neuron cell bodies and along the neurites (Figure 6F). These data strongly suggest that CD63-GFP-positive keratinocyte-derived exosomes are retrogradely transported from the epidermis to DRG neuron cell bodies, representing a novel mode of communication.

**Figure 6:**
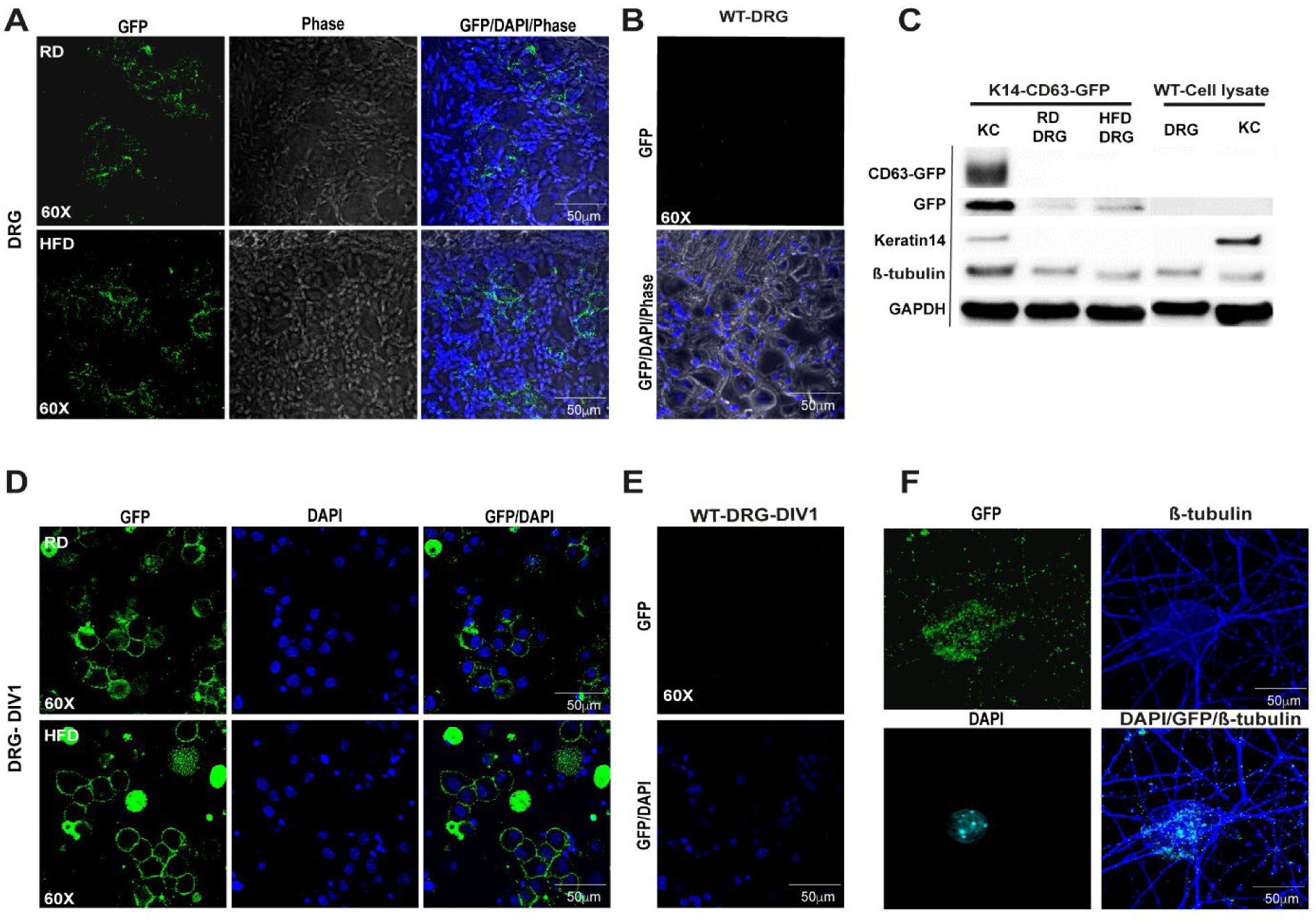
Epidermal keratinocyte-derived exosomes (KDEs) are retrogradely trafficked from the epidermis to DRG neurons *in vivo* in male and female mice. **A)** We detected GFP signal after immunolabel amplification in cryosections of the DRGs from the EV-reporter mice for both RD and HFD. N=3 male biological replicates for both RD and HFD and n=2 for RD female mice. **B)** As expected, no false GFP signal was detect in WT DRG cryosections. N=3 biological replicates of WT. **C)** Immunoblotting the DRG cell lysate of EV-reporter mice revealed GFP expression. N=2 biological replicates for both RD and HFD. **D)** We detected GFP signal after IHC amplification in primary DRG cell cultures from the EV-reporter mice for both RD and HFD. N=3 biological replicates for both RD and HFD. **E)** As expected, no false GFP signal was detected in WT primary DRG cultures. N=3 biological replicates of WT. **F)** The GFP signal was not only detected in the cell body of the primary DRG cultures from EV-reporter mice but also along the neurites, further providing evidence that KDEs are trafficked along the neurites. N=6 biological replicates between RD and HFD.

## DISCUSSION

We isolated and characterized keratinocyte-derived exosomes (KDEs) morphologically and molecularly using an unbiased proteomic and small RNA sequencing approach. Our research revealed that KDEs alter their cargo in a mouse model of painful diabetic neuropathy (PDN), and we identified several gene ontology enrichment pathways that were differentially expressed. In both “omic” datasets, neuron-keratinocyte pathways were enriched both under normal physiology and in PDN. Additionally, we created a K14-CD63-emGFP extracellular vesicle (EV)-reporter mouse line and observed a GFP signal in the DRG neurons and neurites, suggesting that keratinocyte-derived GFP-tagged exosomes are retrogradely trafficked from epidermal keratinocytes into the DRG neurons of mice.

These findings suggest a direct communication pathway between the epidermis and the peripheral nervous system, which is altered in our HFD mouse model of PDN. In our microfluidic paradigm, the presence of keratinocytes enhanced neurite outgrowth, indicating release of a soluble factor by keratinocytes that encourages neurite growth (Figure 2A-C). Proteomic analysis of our KDEs revealed that they contained several catenin and plexin isoforms (3C; Insert). The canonical WNT-signaling pathway plays a significant role in axon development in the central^65,66^ and peripheral^67^ nervous systems and has been implicated in Schwann cell-axon communication^68^. Plexins, the surface receptors for semaphorins, are involved in neuron axonal growth and guidance, as well as a host of other functions^69–72^. Semaphorin 4C-Plexin-b2 signaling has been reported to be markedly increased in states of persistent pain in mice, and downregulation of this pathway led to the impairment of inflammatory hypersensitivity via RhoA-ROCK-dependent mechanisms^73^. Additionally, Sema3A and Plexin A were reported to be dysregulated in the spinal cord of a HFD model of PDN^74^. To our knowledge, this is the first report of these proteins being present as KDE cargo. Additionally, notch1, which was recently reported to facilitate neuron-to-neuron communication through EVs in the hippocampus of mice^46^, was detected in our KDEs for all three biological replicates of both diet groups (3C; Insert), suggesting that this mechanism of internalization may also apply to keratinocyte-neuron terminal endings in the skin. Notch signaling is known to play a prominent role in developing neurons, including in axon guidance^75,76^. Pathogenic variants in the Notch ligand Jagged1 were implicated in the development of peripheral neuropathy in two independent families and confirmed in a mouse model^77^. Thus, Notch signaling may represent an unexplored, novel communication pathway between keratinocytes and DRG neuron terminal nerve endings.

The KDEs from both groups contain differentially expressed synaptotagmin, which is an essential component of the presynaptic vesicle release complex that facilitates vesicle fusion with the plasma membrane^78,79^, along with synaptophysin and synapsin-2, both of which are involved in regulating vesicle docking to the inner plasma membrane^80–82^ (3G; Right Panel). These proteins have all been previously reported in keratinocytes^83^. However, it is still unclear why these proteins are present as KDE cargo and what their function in keratinocyte-to-neuron communication might be. Further experiments are required to validate all these cargo proteins, both in mice and human KDEs, and to investigate their mechanism of action on nerve terminals.

Our small RNA sequencing dataset aligned with the proteomic data. The top two most abundant small RNAs, let-7 and miR-23 (Figure 4A), were predicted to target several prominent signaling pathways that overlapped with the proteomic dataset, including MAPK signaling and the WNT signaling pathways (Figure 4C). A recent study has indicated that partially inhibiting p38-MAPK activation in a diabetic neuropathy rat model led to anti-hyperalgesic effects, suggesting a significant role for MAPKs in nociception modulation^84^. KDEs from both our groups contained a diverse range of MAP kinases (MAPKs), with mapk1 differentially expressed in our HFD model. Both let-7 and miR-23 have been shown to modulate MAPK activity^85,86^. Additionally, two differentially expressed microRNAs suggested altered keratinocyte-to-neuron communication (Figure 4F-G). An altered expression of miR-24-3p has been reported in cancer^62,63,87,88^ and diabetes^89^. By cross-referencing three separate databases for predicted targets and running a gene ontology enrichment analysis, one predicted pathway for miR-24-3p modulation is plexin-semaphorin activity, and our proteomic dataset suggested plexin-b2 as a cargo protein (Figure 3C). However, the precise mechanism by which these microRNAs regulate axon guidance and potentially contribute to PDN remains unclear. Further studies are necessary to understand how these microRNAs modulate target proteins and pathways, either directly or indirectly.

The DRG transcriptome is substantially altered in PDN^90^, resulting in hyperexcitability of the nociceptive neurons that drive neuropathic pain^49,90^ and presenting a possible druggable system for future therapeutics. In our studies, using an EV-reporter mouse line in which KDEs are labelled with CD63-emGFP (Figure 6A), we detected a GFP signal in the DRG neuron cell bodies and neurites (Figure 6F). This indicates that GFP-containing exosomes originating in the epidermis are retrogradely trafficked into the DRG neuron cell body, where presumably they can initiate transcriptional changes due to their cargo. It should be noted that the GFP species detected, at least from our immunoblotting, was the non-fused form of GFP rather than the CD63-GFP fusion form (Figure 6C), but that soluble GFP was also detected in the epidermis (Figure 5C) and KDE fractions (Figure 5E) in high abundance. It may be that the GFP-tagged exosomes are trafficked to DRG neurons, where they release their transcription-altering cargo, and the fusion protein is subsequently degraded into the soluble GFP that we detected in immunoblotting.

This study enhances our understanding of the communication between keratinocytes and DRG neurons and reveals a new role for KDEs in possibly promoting axonal degeneration, which underlies neuropathic pain in PDN. As many genes show differential expression in DRG neurons in PDN mice compared to control mice^91^, our studies support a novel strategy for treating PDN by focusing on the skin rather than the entire body. One of the challenges with current PDN treatments is their systemic administration and off-target effects^10–12^. Here, we present evidence for a delivery pathway from epidermal keratinocytes to the peripheral nervous system, which could be utilized to develop and deliver improved topical treatments for PDN and other nervous system diseases. Additionally, our investigation into the role of exosome-mediated communication between keratinocytes and DRG neurons has broader implications. Indeed, exosomes hold great promise as novel disease biomarkers, therapeutic agents, and drug delivery systems. This potential extends beyond PDN, laying the groundwork for exploring new avenues in pain and peripheral neuropathy research and treatment.

## METHODS

### Animals

Animals were housed on a 12-hour light/12-hour dark cycle with ad libitum access to food and water. We used the following mouse lines: K14-Cre, homozygous; CD63-emGFP fl, homozygous; K14-Cre::CD63-emGFP fl heterogenous.

### HFD

Mice were fed 42% fat (Envigo TD88137) for 10 weeks as a rodent model of type 2 diabetes. Control mice were fed a regular diet (RD) of 11% fat. After 10 weeks of RD or HFD, a glucose tolerance test was performed as described^49^. A cutoff of (≥140 mg/dl) at 2 SD above the mean for glucose 120 minutes after glucose challenge in WT littermate mice was used as a ‘diabetic’ classification^49^.

### Behavioral testing

von Frey behavioral studies were performed as previously described^49^ with random experimental group assignments and double-blind investigator and endpoint analysis conditions.

### Primary keratinocyte cultures

Glabrous paw skin is dissected from the mouse and incubated in dispase (2.3 mg/ml) overnight. The epidermis is separated from the dermis and incubated in TrypLE Express (10 min, 37C; Gibco 12604-013); the keratinocytes are dislodged using gentle agitation with forceps and then plated on 15cm^2^ plates with 154CF epidermal medium (M154CF500) supplemented with 170ul of 0.2M CaCl_2_ and 5ml of HKGS (S-001-5), which was depleted of EVs following 18 hours of ultracentrifugation at 100,000*g*. Medium change occurs 24 hours after plating and then every 48 hours. Complete cell culture medium and all other reagents have been confirmed to be EV-free prior to use with the keratinocyte cultures.

### Wound healing scratch assay

Primary keratinocyte cultures were grown on 6-well plates (Fisherbrand FB012927) with a 300,000-seeding density and grown to 90% confluency. The tip of an Eppendorf 200ul pipette tip (Fisher 02707409) was used to form a vertical scratch down the center. Cultures were tracked every 24 hours on a Leica 2000 LED microscope and analyzed using ImageJ software to measure the rate of gap closure. One-tail paired t-tests were used on the raw dataset to obtain a p-value between groups for each time point (n=58 RD, n=61 HFD for each time point across three separate biological replicates for each group; significance p ≤ 0.05).

### Exosome isolation with size exclusion columns

Cell-conditioned medium from keratinocyte cultures is collected between culture confluency of 60-90%, is centrifuged at 3000*g* for 30 minutes to remove cell debris and is then concentrated to 500 μl using centrifuge size filters (Pierce Protein Concentrators PES 10K, 88528). The sample is then run through an IZON SEC qEV 35nm column (IZON, ICO-35) as previously reported^43^. Fractions 2-3 are used for downstream applications.

### Western Blot

Fractions 1-10 of 400ul each are concentrated to 20ul using Millipore centrifuge filters (Microcon 30kDa, MRCF0R030) and combined with one volume equivalent of BioRad 2x Lammalenni loading buffer (BioRad 1610737) with 5% BME. Each fraction is run through a 3-15% gradient gel (BioRad 45610840) alongside full keratinocyte cell lysate (+) and a 10K pellet (large EVs), running a BCA assay (Thermo Scientific) to load ∼3 μg of protein per well. Protein was then transferred to PVDF membranes (Millipore) and blocked (BioRad Everyblot 12010020) for 15 minutes. Primary antibodies are applied overnight at 4 °C with secondary antibody at room temperature for 2 hours with 3 TBST washes between each step. Proteins are visualized with a chemiluminescence detection system (Thermo Scientific 32209). All western blot gels were run at minimum in triplicate. Blots were visualized on a Li-Cor Odyssey Fc.

### Western blot antibodies

The following primary antibodies were used: Alix (Abcam ab88388), Syntenin-1 (Abcam ab19903), Tsg101 (Invitrogen PA531260), CD63 (Invitrogen 2H5I1), CD81 (Abcam ab109201), GM130 (Abcam ab52649), Calnexin (Abcam ab22595), K14 (BioLegend 906004), GFP (Abcam ab13970), β-tubulin (ProteinTech 80713-1-RR100UL), GAPDH (Abcam ab181602). The following secondary antibodies were used: Goat anti-rabbit HRP (Abcam ab97080), Goat anti-chicken HRP (Invitrogen A16054).

### Dynamic light scattering

80ul of samples are pipetted into cuvettes (Malvern Catalog#759200) and run through the zetasizer spectrophotometer (Malvern Zetasizer Nano ZSP) with an EV refractive index of 1.35 in ANTEC through Northwestern University. Malvern analytical software is used to analyze the output using particle counts relative to the signal intensity.

### Negative stain EM

Samples suspended in PBS are prepared using the standard uranyl acetate fixation for 5 mins seeded on EMS grids (TMS Catalog#71150) and imaged on a FEI Spirit 2 electron microscope. Images were processed using ImageJ software.

### Cryo-EM

CryoEM images are obtained through the northwestern BioCryo core facility (NUANCE) with samples prepared as previously described^92^.

### Primary DRG Cultures

DRG sensory neurons from WT and K14-CD63-GFP mice were dissociated as described^93^ at 18 weeks of age.

### Microfluidics

Primary DRG cultures were deposited into one compartment chamber connected to the microchannel column in a microfluidic system (XONA Microfluidics SND450) with or without primary keratinocyte cultures in the other compartment chambers. Keratinocytes were cultured for 3 days before depositing DRG cultures unless otherwise specified. The medium was a combination of 50% DRG culture medium as previously described^93^ and 50% keratinocyte culture medium when both cell types were present in the chambers.

### DIR Labelling

Concentrated CCM was labeled with DIR (Invitrogen D12731; 2ug/ul) at a ratio of 1:100 by volume and then passed through the IZON 35nm columns. Isolated DIR-labeled exosomes from fractions 2-3 were then used for downstream applications with a negative DIR control, which was DIR-added to concentrated EV-depleted unconditioned medium passed through the column with the same fractions collected for experiments.

### DIR-Exosomes

DRG Cultures: DIR-labeled exosomes were directly applied to DIV-1 primary DRG cultures and visualized after 16 hours post-treatment by confocal microscopy. Microfluidics: DIR-exosomes were applied to the empty compartment after DRG neuron neurites occupied all microchannels, preventing medium exchange between the two compartments. The microfluidic chambers were visualized after 16 hours by confocal microscopy.

### MS sample preparation

Trichloroacetic acid (TCA, Sigma-Aldrich, Cat# T0699) precipitation was used to clean and precipitate proteins from EV samples. Protein pellets were resuspended in 8 M urea (ThermoFisher Scientific, Cat # 29700) prepared in 100 mM ammonium bicarbonate solution (Fluka, Cat # 09830) and processed with ProteaseMAX (Promega, Cat # V2072) according to the manufacturer’s protocol. The samples were reduced with 5 mM Tris(2-carboxyethyl)phosphine (TCEP, Sigma-Aldrich, Cat # C4706; vortexed for 1 hour at RT), alkylated in the dark with 10 mM iodoacetamide (IAA, Sigma-Aldrich, Cat # I1149; 20 min at RT), diluted with 100 mM ABC, and quenched with 25 mM TCEP. Samples were diluted with 100 mM ammonium bicarbonate solution, and digested with Trypsin (1:50, Promega, Cat # V5280) for overnight incubation at 37°C with intensive agitation. The next day, reaction was quenched by adding 1% trifluoroacetic acid (TFA, Fisher Scientific, O4902-100). The samples were desalted using Peptide Desalting Spin Columns (Thermo Fisher Scientific, Cat # 89882). All samples were vacuum centrifuged to dry.

### Tandem Mass spectrometry

Three micrograms of each sample were auto-sampler loaded with a Thermo Vanquish Neo UHPLC system onto a PepMap™ Neo Trap Cartridge (Thermo Fisher Scientific, Cat#: 174500, diameter, 300 µm, length, 5 mm, particle size, 5 μm, pore size, 100 Å, stationary phase, C18) coupled to a nanoViper analytical column (Thermo Fisher Scientific, Cat#: 164570, diameter, 0.075 mm, length, 500 mm, particle size, 3 µm, pore size, 100 Å, stationary phase, C18) with stainless steel emitter tip assembled on the Nanospray Flex Ion Source with a spray voltage of 2000 V. An Orbitrap Ascend (Thermo Fisher Scientific) was used to acquire all the MS spectral data. Buffer A contained 99.9% H2O and 0.1% FA, and buffer B contained 80.0% ACN, 19.9% H2O with 0.1% FA. For each fraction, the chromatographic run was for 2 hours in total with the following profile: 0-8% for 6, 8% for 64, 24% for 20, 36% for 10, 55% for 10, 95% for 10 and again 95% for 6 We used Orbitrap HCD-MS2 method for these experiments. Briefly, ion transfer tube temp = 275 °C, Easy-IC internal mass calibration, default charge state = 2 and cycle time = 3 s. Detector type set to Orbitrap, with 60K resolution, with wide quad isolation, mass range = normal, scan range = 375-1500 m/z, max injection time mode = Auto, AGC target = Standard, microscans = 1, S-lens RF level = 60, without source fragmentation, and datatype = Profile. MIPS was set as on, included charge states = 2-7 (reject unassigned). Dynamic exclusion enabled with n = 1 for 60s exclusion duration at 10 ppm for high and low with Exclude Isotopes. Isolation Mode = Quadrupole, isolation window = 1.6, isolation Offset = Off, active type = HCD, collision energy mode = Fixed, HCD collision energy type = Normalized, HCD collision energy = 25%, detector type = Orbitrap, orbitrap resolution = 15K, mass range = Normal, scan range mode = Auto, max injection time mode = Auto, AGC target = Standard, Microscans = 1, data type = Centroid.mins receptively.

### MS data analysis and quantification

Protein identification/quantification and analysis were performed with Integrated Proteomics Pipeline - IP2 (Bruker, Madison, WI. http://www.integratedproteomics.com/) using ProLuCID^94,95^, DTASelect2^96,97^, Census and Quantitative Analysis. Spectrum raw files were extracted into MS1, MS2 files using RawConverter (http://fields.scripps.edu/downloads.php). The tandem mass spectra (raw files from the same sample were searched together) were searched against UniProt mouse (downloaded on 07-29-2023) protein databases^98^ and matched to sequences using the ProLuCID/SEQUEST algorithm (ProLuCID version 3.1) with 50 ppm peptide mass tolerance for precursor ions and 600 ppm for fragment ions. The search space included all fully and half-tryptic peptide candidates within the mass tolerance window with no-miscleavage constraint, assembled, and filtered with DTASelect2 through IP2. To estimate protein probabilities and false-discovery rates (FDR) accurately, we used a target/decoy database containing the reversed sequences of all the proteins appended to the target database^98^ (UniProt, 2015). Each protein identified was required to have a minimum of one peptide of minimal length of six amino acid residues. After the peptide/spectrum matches were filtered, we estimated that the protein FDRs were ≤ 1% for each sample analysis. Resulting protein lists include subset proteins to allow for consideration of all possible protein isoforms implicated by at least three given peptides identified from the complex protein mixtures. Then, we used Census and Quantitative Analysis in IP2 for protein quantification. Static modification: 57.02146 C for carbamidomethylation. Quantification was performed by the built-in module in IP2.

### Proteomics Statistics

The spectral counts for each protein accession ID for the three RD and HFD biological replicates are used to run a one-tailed, paired t-test for statistical significance. The average spectral counts for RD and HFD were used to obtain a HFD/RD fold change. The volcano plot was generated in R using ggplot2. The heat map was generated using the R package pheatmap. Gene ontology enrichment analyses were generated using the R packages clusterprofiler, msigdbr and ggplot2. GO enrichment analyses used proteins present in all biological replicates of each group only.

### Small RNA sequencing

Samples underwent small RNA sequencing through Northwestern’s core facilities. Biological triplicate RD and HFD RNA samples were prepared using the total exosome RNA and protein extraction kit from Invitrogen (Invitrogen Catalog#2743605). RNA samples were quantified by Qubit RNA HS assay and the quality was confirmed by Bioanalyzer RNA pico chip assay. Then, 1ng of RNA was used as input for library preparation with NextFlex small RNA-seq kit v4 according to manufacturer’s protocol. Each sample was barcoded with a unique index and multiplexed libraries were pooled for sequencing on Novaseq X Plus 10B flowcell using single end 50nt mode.

### RNA sequencing Statistics

Data analysis was carried out in R using the standard workflow of DESeq2 paired with the libraries apeglm and ggplot2 for figure generation. Bonferroni post-hoc adjustment was used for the reported adjusted p-values ≤ 0.05 for significance. Gene ontology enrichment analyses were generated using the R packages clusterprofiler, msigdbr and ggplot2 with Bonferroni post-hoc adjusted P values < 0.05 for significance.

### Protein target analysis

The direct or indirect protein targets for each microRNA was predicted using several target prediction programs, including miRDB (http://mirdb.org/)^99^, TargetScan v7.0 (http://www.targetscan.org/vert_72/) and DIANA-microT v5.0 (https://bio.tools/DIANA-microT). Only the predicted proteins identified by all 3 programs were included in the subsequent enrichment analyses.

### Immunohistochemistry

Glabrous hind paw dermis/epidermis was separated from the paw and whole DRGs (lumber 2-4) were isolated from 18 week old mice and fixed with 4% PFA for 1 hour, 30% sucrose for 1 hour, and then embedded in OCT. Samples were processed as previously described^50^ and analyzed by confocal microscopy.

### Antibodies

We used the following primary antibodies on DRG sections: GFP (chicken; Abcam ab13970). We used the following antibodies on skin sections: K14 (BioLegend 906004). We used the following antibodies on primary DRG cultures: GFP (Abcam ab13970), β-tubulin (ProteinTech 80713-1-RR100UL). We used the following secondary antibodies: Goat anti-chicken AlexaFluor^TM^-598 (Invitrogen A-11042), goat anti-rabbit AlexaFluor-647 (Invitrogen A32733).

### EV-Reporter Mouse

The commercially available CD63-GFP fl/fl mouse line (Jackson Laboratory Strain#:036865) was crossed with K14-Cre mice to generate the EV-reporter K14-CD63-GFP mouse line.

### Study approval

All methods involving animals were approved by the IACUC of Northwestern University.

## ACKNOWLEDGEMENTS

Zetasizer Nano ZSP (DLS), Nanosight (NTA), and IZON Exoid (TRPS) experiments were performed in the Analytical bioNanoTechnology Equipment Core Facility of the Simpson Querrey Institute for BioNanotechnology at Northwestern University. ANTEC receives partial support from the Soft and Hybrid Nanotechnology Experimental (SHyNE) Resource (NSF ECCS-2025633) and Feinberg School of Medicine, Northwestern University. Electron microscopy imaging work was performed at the Northwestern University Center for Advanced Microscopy (RRID: SCR_020996) generously supported by NCI CCSG P30 CA060553 awarded to the Robert H Lurie Comprehensive Cancer Center. This work made use of the EPIC facility of Northwestern University’s NUANCE Center, which has received support from the SHyNE Resource (NSF ECCS-2025633), the IIN, and Northwestern’s MRSEC program (NSF DMR-2308691). And this work was supported by the Northwestern University NUSeq Core Facility.

## STATEMENT OF DATA AND MATERIALS

We will make the raw MS data publicly available in an accessible database upon acceptance.

