## Supplemental Figures for "Keratinocyte-Derived Exosomes in Painful Diabetic Neuropathy"

### SUPPLEMENTARY FIGURES

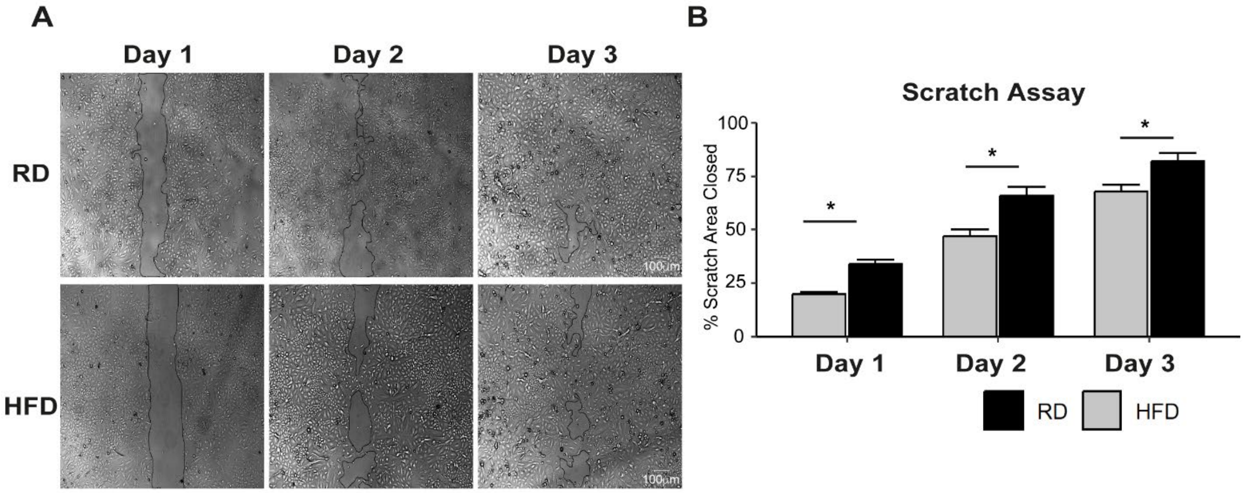


**Supplemental Figure 1: Keratinocytes from HFD mice exhibit impaired wound healing in primary cultures. A)** HFD keratinocytes exhibit impaired wound healing compared to their RD controls (n=58 RD, 61 HFD over 3 biological replicates for each day). **B)** Quantification of the scratch assay (one-tail, paired t-test; $p \leq 0.05$ for significance). Bars are mean ± SEM.

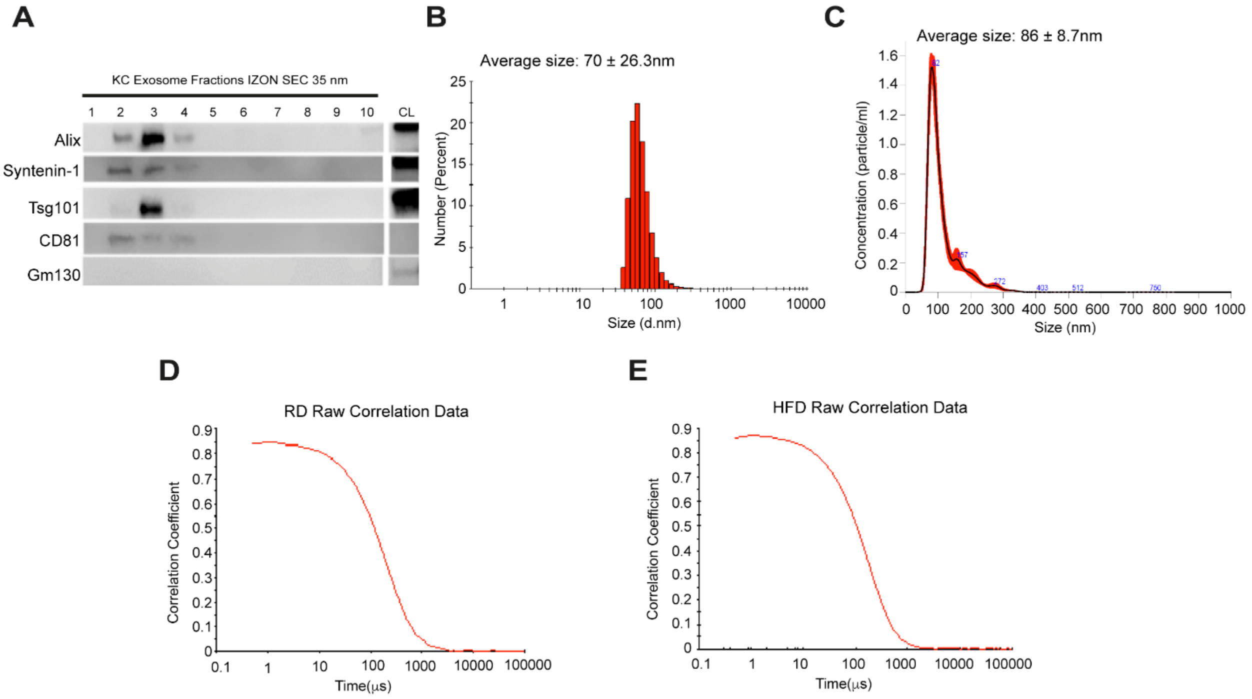


**Supplemental Figure 2: Keratinocyte-derived exosome (KDE) characterization for HFD and DLS quality controls. A)** Fractionated cell-conditioned medium from HFD keratinocyte cultures revealed exosome-associated proteins enriched in SEC fractions 2/3. **B)** SEC Fr2/3 for HFD KDEs had an average size range of 70 ± 26.3nm (mean ± StDev) with dynamic light scattering. N=3 biological replicates from 2 male mice and 1 female mouse. **C)** KDEs analyzed using nanoparticle tracking analysis produced a concentration peak particle size of 86 ± 8.7nm (mean ± StDev). N=2 male biological replicates. **D)** Raw correlation curve for RD DLS KDEs dataset passes quality control. **E)** Raw correlation curve for HFD DLS KDEs dataset passes quality control.

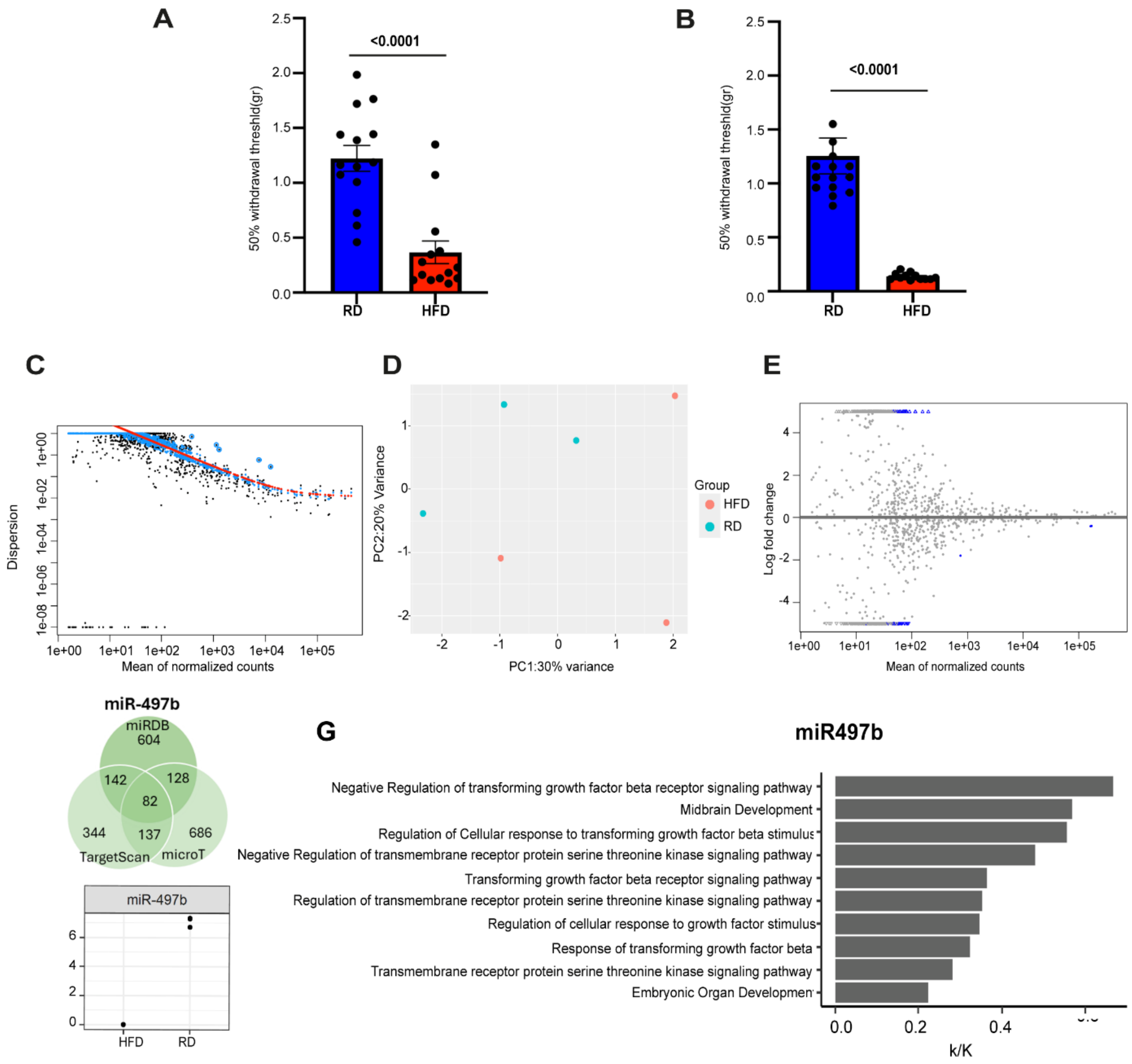


**Supplemental Figure 3: The proteomic and RNAsequencing datasets represent painful diabetic neuropathy. A)** Male mice used in the proteomic experiment were tested for mechanical allodynia. HFD mice after 10 weeks of diet presented with a lower withdrawal threshold. N=12 biological replicates for each group. P values were calculated by two-tail Mann-Whitney test. **B)** Mice used in the RNAsequencing experiment were tested for mechanical allodynia. HFD mice after 10 weeks of diet presented with a lower withdrawal threshold. N=12 biological replicates for each group. P values were calculated by two-tail Mann-Whitney test. **C)** As expected, the dispersion of small RNA reads decreased with increasing mean normalized counts. **D)** Biological replicates of RD and HFD acceptably grouped together based on variance in the PCA plot. **E)** Small RNAs identified in high concentration relative to the whole experienced decreased fold change differences between the two groups as expected. The one significantly differentially expressed small RNA with a high mean count was miR-24-3p, represented as the far right blue datapoint. **F)** We found that miR-497b was downregulated in HFD and used three separate databases for predicted protein targets. **G)** The gene ontology enrichment analysis for these predicted proteins presented several interesting predicted GO Term pathways, including the TGFβ signaling pathway. GO Terms Bonferroni adjusted p-value $\leq 0.01$.

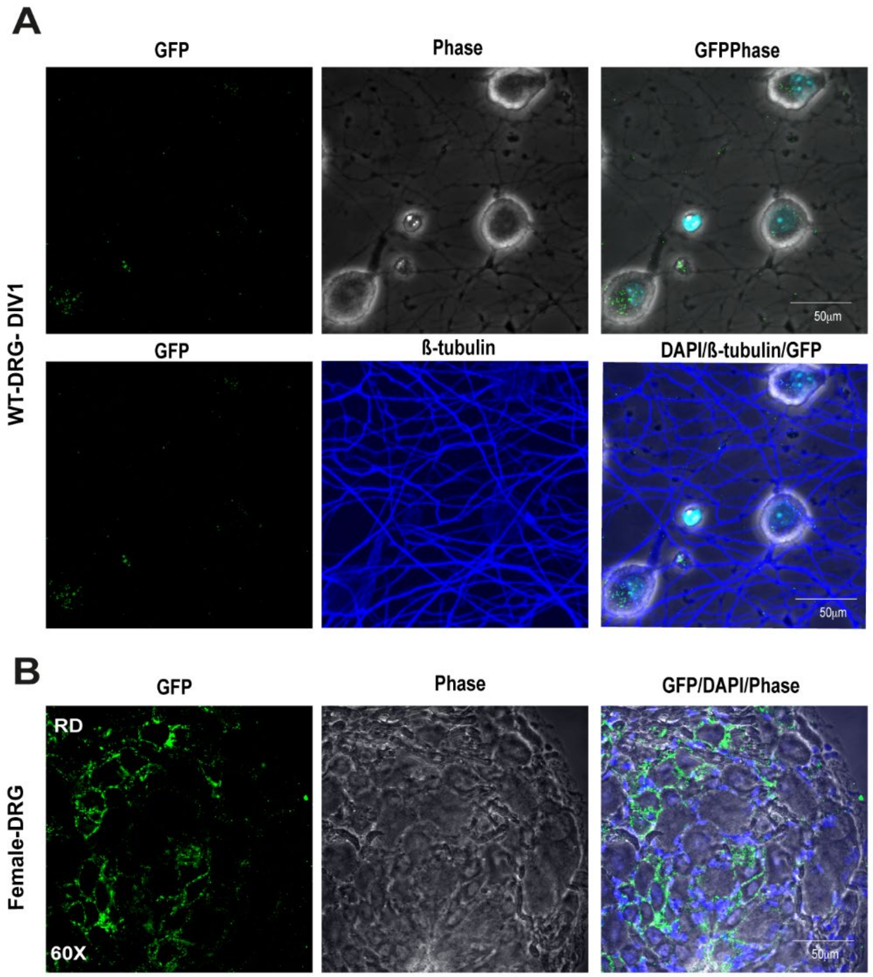


**Supplemental Figure 4: GFP-tagged KDEs are internalized by DRG neurons *in vitro*. A)** KDEs from SEC Fr2/3 of EV-reporter keratinocyte cell conditioned medium, whereby the KDEs are GFP-tagged, are functionally internalized by primary WT DRG neurons and observed in both the cell body and neurite with either phase (top panels) or β-tubulin co-staining (bottom panels). N=4 across 2 biological replicates. **B)** We confirmed GFP signal after immunolabeling amplification in cryosections of the DRGs from EV-reporter female RD mice. N=4 across 2 female biological replicates.
